# *FPtool* a software tool to obtain *in silico* genotype-phenotype signatures and fingerprints based on massive model simulations

**DOI:** 10.1101/266775

**Authors:** Guido Santos, Julio Vera

**Author notes:** Corresponding author: Prof. Dr. Julio Vera, Universitätsklinikum Erlangen, Hartmanssstr. 14, Erlangen, Germany.

## Abstract

*Fptool* is an intuitive tool that provides to the user a preliminary fingerprint of the behaviour simulated by a mathematical model of a biochemical network when comparing two biological scenarios defined by the user. Here we present the tool and we applied to an already published mathematical model of lung legionella infection. The fingerprint obtained correlates with the results obtained in the original article. This tool is optimal for the users that would like to obtain a fast and preliminary view of the qualitative behaviour of a mathematical model before deciding for more elaborate analyses.

## Introduction

Mathematical modeling based simulation of biological systems helps us to understand the complexity of molecular, cellular and individual interactions^1^. Giving the limitations for producing sufficient amount of reliable quantitative data, the main drawback of this approach is the calibration of the models to obtain accurate values for the model parameter^2,3^. An alternative approach to parameter estimation is to use computational analysis and systematic simulations to analyse the whole structure of the model either analytically^4^ or numerically^5^. By doing this one can obtain a qualitative view of the different biological behaviours that the model can generate and are encoded as subspaces of model solutions. The analysis performed is a type of global sensitivity analysis. Here we propose a tool to obtain a preliminary view of the qualitative behaviour of a mathematical model, referred as fingerprint.

## Methods

The main aim of *FPtool* (*FingerPrint tool*) is obtaining a fingerprint of two different biological scenarios defined by the user. The mathematical model should be encoded in a way that it accepts a vector of parameters (input vector) as input and it produces as an output either a 1 or a 0 depending to the scenario that this solution belongs. The user code of the model must be written in a way it performs model simulations for the input vector and decides to which scenario the input vector is assigned based in output of the simulation. The tool generates a user-defined number of random model solutions, classify them based on the output defined and taken all of them as an ensemble of model simulations, generate a fingerprint based on the distribution of the parameters values in the input vectors for each of the two biological scenarios compared underlying the in each scenario. Previous examples for the application of this strategy are contained in Vera et al. 2013, Santos et al. 2016 and Nikolov et al. 2018^6–8^.

The default analysis produces a figure with the distribution of parameter values in both scenarios (upper panels in Figure 1). This is the first fingerprint that can be used to compare the scenarios by the user. Furthermore, the user can decide to select a subset of the parameters to perform two cluster analyses on the different scenarios (lower panels in Figure 1). From this cluster analysis it can be compared the similarity between the parameters depending on the biological scenario defined. In the Application section the tool is applied to a published model to obtain the fingerprint in two different scenarios (Figure 1). This example model is included as example in the *PFtool* package.

The current version of the tool runs on Matlab (tested on Matlab 2015b 64 bits) and it accepts Matlab scripts as input. The tool produce as an output csv files with the groups of solutions. I second, sequential step, the files produced in the analysis can be used in R (version 3.4.3) to generate a deeper analysis of the solutions, as indicated in the instructions provided in the *Readme.txt* file. The R-based analysis implements a logistic regression analysis in R of the data retrieved in the simulations.

## Application

We used the tool with the model presented in Schulz et al. 2017^9^. Precisely, we used the mathematical model proposed by the authors to obtain a fingerprint between two different biological scenarios. The model simulates the activation of the NFkB intracellular signalling pathway and the production of IL-8 in lung epithelial cells infected with the Legionella pathogen. The two scenarios defined relied on the amount of IL-8 simulated by the model after 50 hours infection. If the amount of IL-8 is higher than 24 ng/ml the solution will be part of the scenario 1, while if it is lower it will belong to the scenario 0. The input vector includes the values of the 24 model parameters. Figure 1 shows the fingerprint of both scenarios in the upper panels. The distribution of values of parameters can be compared to investigate the differences between the in the scenarios. From left to right there are presented the parameters with higher difference between the scenarios. The lower panels present the second fingerprint based on cluster analysis. It clusters the parameters based on their correlation between different solutions.

**Figure 1.**
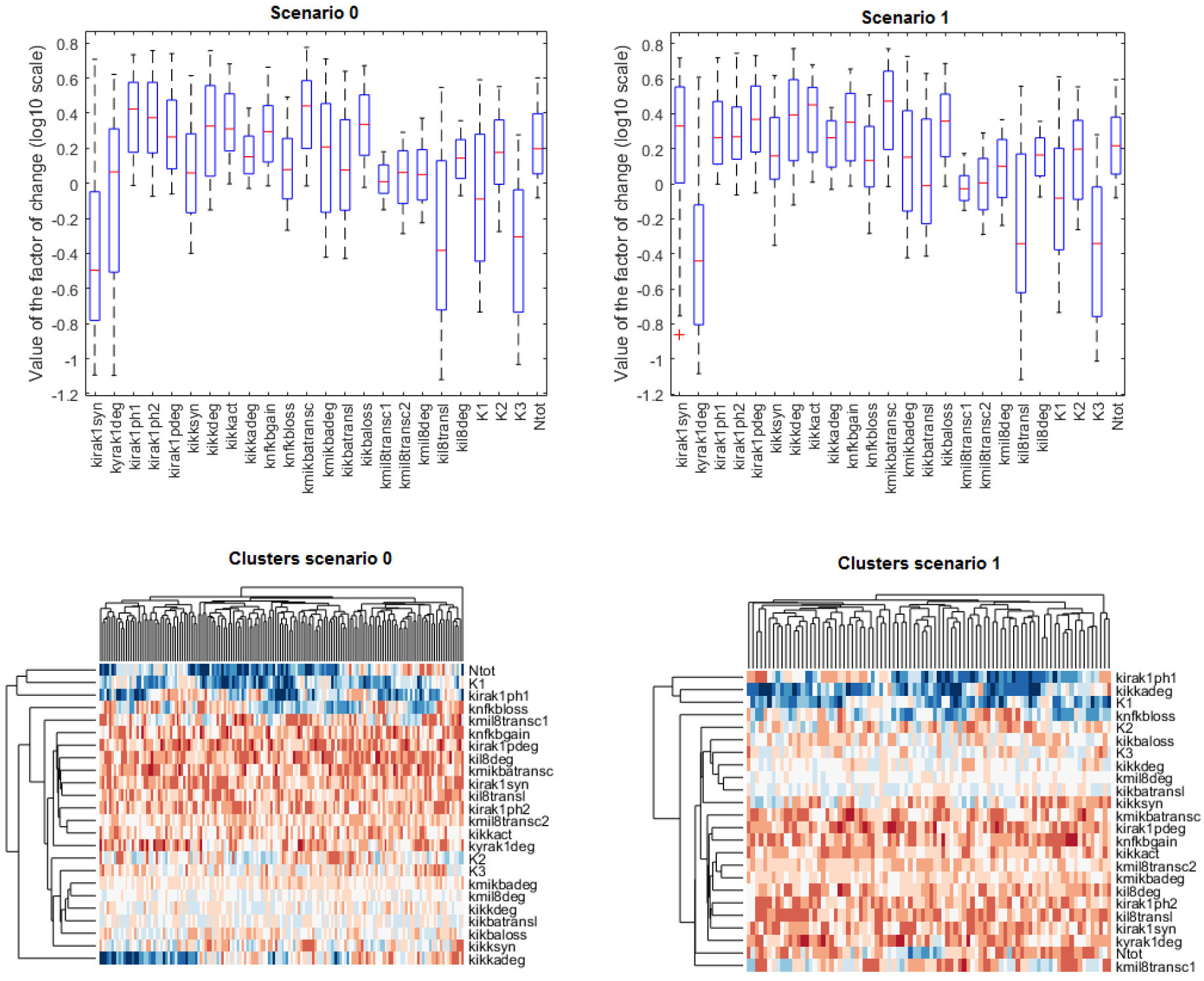
Fingerprints of the model taken from Schulz et al. 2017^9^ obtained by *FPtool*.

The fingerprint in the upper figures points to the Irak1 related parameters as the ones that differentiate the biological scenarios the best. This fingerprint correlates with the results of the original article, in which the authors propose Irak1 one of the leading factors controlling the IL-8 production^9^. The lower figures extend the fingerprint by clustering the input vectors in sub-classes within the two output scenarios considered in the analysis.

## Conclusion

*FPtool* provides to the user a general idea of the qualitative behaviour of a mathematical model by obtaining a fingerprint of two different biological scenarios. The information obtained by this tool can help the user to identify potential parameters that segregate the biological scenarios of interest.

## Availability

The tool can be downloaded at https://zenodo.org/record/1173896#.WoaUHK7iaUk

